# Xylem traits and growth phenology predict growth and mortality response to defoliation in temperate forests

**DOI:** 10.1101/030015

**Authors:** Jane R. Foster

## Abstract

Defoliation outbreaks are biological disturbances that alter tree growth and mortality in temperate forests. Trees respond to defoliation in many ways; some recover rapidly, while others decline gradually or die. These differences may arise from species functional traits that constrain growth such as xylem anatomy, growth phenology or non-structural carbohydrate (NSC) storage, but this has not been shown. Although many studies address these phenomena, varied and idiosyncratic measures limit our ability to generalize and predict defoliation responses across species. I synthesized and translated published growth and mortality data into consistent standardized variables suitable for numerical models. I analyzed data from 32 studies, including 16 tree species and 10 defoliator systems from North America and Eurasia, and quantitatively compared responses to defoliation among species and tree functional groups using linear mixed-effects models.

Relative growth decreased linearly or curvilinearly as defoliation stress accumulated across species. Growth decreased by only 5-20% following 100% defoliation in ring-porous *Quercus,* whereas growth of diffuse-porous hardwoods and conifers declined by 50-100%. Mortality increased exponentially with defoliation, more rapidly for *Pinus* and diffuse-porous species than for *Quercus* and *Abies.* Species-specific mixed models were best (R^2^c = 0.83-0.94), yet functional-group models lost little in terms of goodness-of-fit (R^2^c = 0.72-0.92), providing useful alternatives when species data is lacking. These responses are consistent with functional differences in wood growth phenology and NSC storage. Ring-porous spring xylem growth precedes budburst. Defoliators whose damage follows foliar development can only affect development of later wood. Growth of diffuse-porous and coniferous species responds more drastically, yet differences in NSC storage make them more vulnerable to mortality as stress accumulates. Ring-porous species resist defoliation-related changes in growth and mortality more than diffuse-porous and coniferous species. These findings apply in general to disturbances that cause spring defoliation and should be incorporated into forest vegetation models.

## Introduction

Tree and forest response to spring defoliation is a complex process. Trees lose photosynthetic potential when they are defoliated early in the growing season, resulting in changes in growth. They also die more often as defoliation stress interacts with other mortality factors. Yet trees are long-lived, sessile organisms that must weather many disturbances over their lifetimes. In adapting to this life history, they have become resilient to a wide range of climatic variation and disturbance, including some amount of herbivory and defoliation. This adaptive resilience can mask the effects of defoliation when disturbance years are observed in isolation, as some trees show negligible response. Yet as stress accumulates over successive years of an outbreak, changes in tree growth and mortality become more severe, consistent and interpretable (MacLean 1980).

Tree responses to stress are increasingly viewed through the lens of inter-specific differences in functional traits including leaf and wood growth phenology, xylem anatomy, and non-structural carbohydrate (NSC) storage (Wiley & Helliker 2012, Panchen et al. 2014, Sevanto et al. 2014). Ensembles of these traits are often linked. For example, deciduous species that leaf out later tend to have larger xylem vessels than species that leaf out earlier (Lechowicz 1984, Panchen et al. 2014), while species that store NSC primarily in leaves (conifers) tend to break bud latest of all (Hoch et al. 2003, Michelot et al. 2012). These linkages may help explain mechanisms of tree mortality under drought or other stresses (Sevanto et al. 2014), yet a consensus on mortality mechanisms remains elusive (Wiley & Helliker 2012). Insect defoliation is a periodic stressor of trees that is rarely brought to bear on ongoing discussions of carbon starvation or hydraulic limitation hypotheses (Anderegg & Callaway 2012, Landhäusser & Lieffers 2012). Yet defoliation stress is relatively straightforward: it directly limits the mainsource of C during a critical growing period. A better understanding of functional differences in tree response to defoliation would provide insight into responses to distinct climate-related stresses such as drought or late frost.

Extensive research quantifies growth and mortality responses to defoliation, yet it remains difficult to generalize the results across species and ecosystems due to the wide variety of data, models, and scales used by individual studies (Feicht et al. 1993, Hallet et al. 2006). These disparities present a significant obstacle to the development of general models that simulate how defoliation affects forest productivity. While extensive reviews exist for general (Kulman 1971) and species-specific defoliation effects (Davidson et al. 1999; Jacquet et al. 2012), they stop short of quantitatively synthesizing results to compare defoliator systems (but see MacLean 1980). Comparisons are complicated by the common use of categorical measures of defoliation that differ. These measures may be defined by quantitative limits (e.g. low, medium, and high defoliation classes may correspond to 0-30%, 30-60%, 60-100% defoliation, or other limits), but they are typically reported and discussed categorically. These limitations leave a gap in our understanding about forest response to large-scale, recurring insect disturbance that contributes significant uncertainty to landscape-and global vegetation models.

We can reduce uncertainty in forest models by quantifying the different ways species respond to stress and how those responses are linked to functional traits. We can also improve how observations from field-based studies are scaled to geographic extents that are compatible with vegetation models. Recent innovations using satellite data to quantify and map defoliation create the potential to link spatially accurate estimates of disturbance stress with forest characteristics and response (Townsend et al. 2012, Foster et al. 2013). In order to improve forest models with realistic representations of defoliation severity, we need numerical relationships that link accumulated defoliation stress to growth suppression or mortality. Examples of such empirical relationships exist for few defoliator systems (MacLean 1980; Alfaro et al. 1982). In this analysis, I compare defoliation responses among several species from different tree functional groups. I converted published data to standardized variables that lend themselves to stand- and landscape-scale forest models and examined the data for significant trends, relationships and differences among defoliator systems. Specifically, I sought studies that allow quantification of both accumulated defoliation stress, which I define in a manner similar to MacLean (1980) as the sum of annual defoliation over multiple years of an outbreak, and the associated responses in terms of growth and mortality. I expected sensitivity to defoliation to differ both by species and by functional groups defined by general tree-growth strategies.

## Materials and methods

To test for generalizable relationships, I compiled research papers that reported defoliation as a percentage of whole canopy foliage, or alternatively cumulative defoliation (defined as the sum of annual defoliation over an outbreak or the product of average defoliation and outbreak duration), as well as changes in growth and mortality as percentages (growth = % of average or expected growth, cumulative mortality = % of population dying over a period of time, typically 5-10 years). Because repeated defoliation causes cumulative stress in trees, data from multiple years of defoliation are of particular interest when modeling long-term effects (Blaise 1958; MacLean 1980; Colbert & Fekedulegn 2001; Hennigar, MacLean & Norfolk 2007). Increasing intensity over the duration of an outbreak can be captured by the summed, cumulative defoliation in an objective, quantitative way (MacLean & Ostaff 1989). For papers where data were presented in tables, graphs or text that could be converted to the described variables, I extracted, transformed and analyzed them in a consistent manner. I assigned categorical annual or cumulative defoliation values to the numerical midpoint of the reported defoliation range (e.g. a 150-300% cumulative defoliation category, became 225% defoliation) (MacLean & Lidstone 1982). I treated the spruce budworm system (SBW) (*Choristoneura fumiferana*) as a special case because cumulative defoliation is defined uniquely in this literature as an index of defoliation by foliage age-classes. To retain comparison with existing published relationships, I did not rescale these data to fit my definition of cumulative “total” canopy defoliation (e.g. simultaneous defoliation of all foliage age-classes).

I selected results that reported the aggregate response of all tree canopy classes when available, as many forest models do not track individual trees or their canopy status (Mladenoff 2004). When data were only reported for separate canopy classes (i.e. suppressed, intermediate, or dominant) and sample sizes were available, I used weighted averages of all classes to estimate aggregate stand response (Campbell & Valentine 1972). I did not distinguish among differences in growth or mortality variables, such as mortality expressed as a percentage of basal area (BA) or number of stems. My goal was to determine if average trends could be found in the extensive empirical literature and whether trends would vary among tree species groups. I expected considerable variability in both the predictor and response variables across studies, as the methods of their estimation often lacked precision. Many papers only reported means of cumulative defoliation categories, and relationships fit to mean values mask the variability present in the raw, unreported data. The R^2^ values reported from mean-only data do not represent the variability expected from plot-level data, which would presumably result in lower R^2^.

I fit mortality response (%) to cumulative defoliation using a log transform to linearize the relationship. I fit either linear or negative log-linear relationships between cumulative defoliation and growth (%) where appropriate. I report parameters and results in tables and back-transformed models in figures with the raw data. When possible, I also fit models to annual mortality rates, calculated by dividing raw reported mortality by the number of years over which it had accumulated. When multiple datasets for similar species or defoliator systems existed, I used mixed models (nlme package in R) to create combined models for annual rates that varied intercept and slope as random effects by study and compared the results using Akaike information criterion for small sample sizes (AICc).

## Results

### Growth Response

Average radial growth decreased linearly with accumulated defoliation stress for 9 of 16 data sets that reported usable data, and followed a negative exponential relationship for the remainder (Table 1, Figs. 1 and 2). Cumulative defoliation explained from 17 to 99% of the variance in relative growth depending on the dataset, with typical R^2^ values exceeding 0.80 (Table 1). The rate of growth suppression, i.e., the slope parameter, tended to vary bimodally. More sensitive tree species-defoliator systems exhibited a negative slope equal to a 50-100% suppression in growth for a 100% increase in cumulative defoliation (Fig. 2a) (Kulman, Hodson & Duncan 1963; Alfaro & Shepherd 1991; Gross 1992; Erdle & MacLean 1999), while growth of other species was reduced by only 5-20% over the same defoliation range (Figs. 1, 2 and 3, Table 1, Appendix S1 in Supporting Information) (Baker 1941; Rose 1958; Rubtsov 1996; Colbert & Fekedulegn 2001; Naidoo & Lechowicz 2001). Conifers including *Abies balsamea, Pinus banksiana, P. pinaster, P. sylvestrus*, *Psuedotsuga menziesii*, *Larix laricina*, and diffuse-porous *Populus* and *Acer* species composed the more responsive group, showing immediate, strong responses to a single year of defoliation. Species with slower, more moderate growth response generated some radial growth even under severe defoliation, a trait characteristic of ring-porous *Quercus* species, which have positive minimum growth thresholds due to their need to grow new functional xylem cells every spring (Fig. 1). Trees with growth falling below these thresholds would presumably reach a point from which they are unable to recover and subsequently leave the pool of survivors. The lone deciduous conifer, *L. laricina,* responded less rapidly than *Pinus* species, reducing growth at rates intermediate between examples of *P. tremuloides* and juvenile *A. saccharum* (Appendix S1).

**Figure 1.**
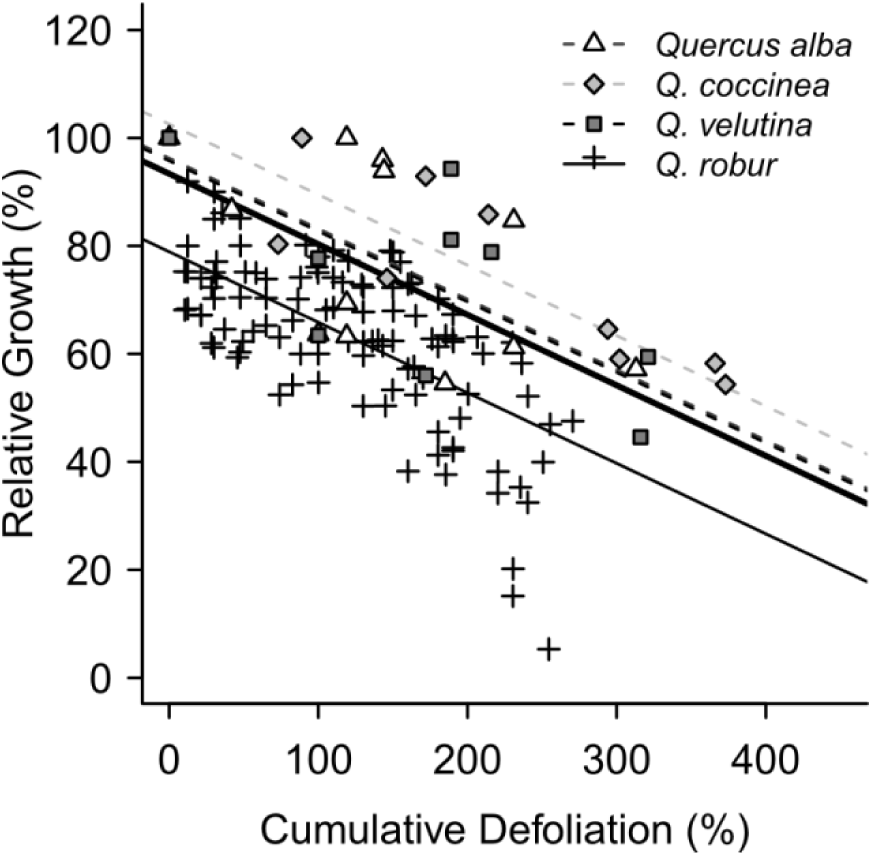
Growth response of *Quercus* species to *Lymantria dispar* (L.) (gypsy moth) defoliation including *Quercus robur* (Rubstov 1996), *Q. velutina* (Baker 1941), *Q. rubra* (Wargo 1981), and *Quercus alba* (Baker 1941, Wargo 1981). Lines show best fit mixed model (Appendix S3, Table S3-1). The best combined *Quercus* model varied intercepts but not slopes.

**Figure 2.**
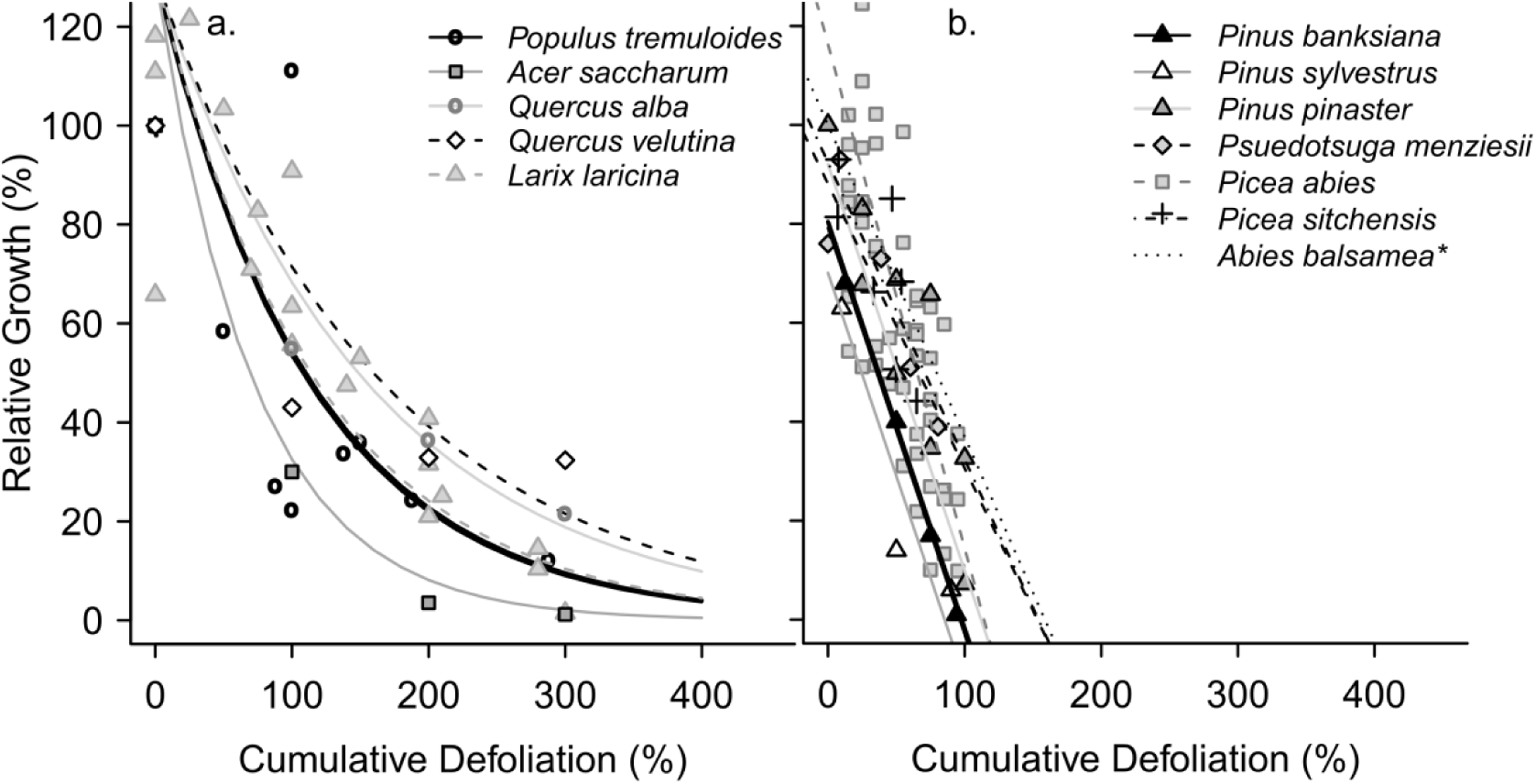
Growth decreases curvilinearly to defoliation (a.) in *Populus tremuloides* (Rose 1958), manually defoliated juvenile trees of *Acer saccharum, Q. alba,* and *Q. velutina* (Wargo 1981), and *Larix laricina* (Ives and Nairn 1966). Linear growth responses of coniferous species (b.) include steep declines in *Pinus banksiana*, *P. pinaster*, *P. sylvestrus* with slower declines in *Pseudotsuga menziesii, Picea abies,* and *Abies balsamea.* The best fit mixed model (a., lines) randomly varied slopes but not intercepts (Appendix S3, Table S3-1). Lines in (b.) show model fits from Table 1 and the best mixed model for *Pinus* species that varied intercept, but not slope (Appendix). *A. balsamea* relationship was published by Dobesberger 1998.

**Table 1.**
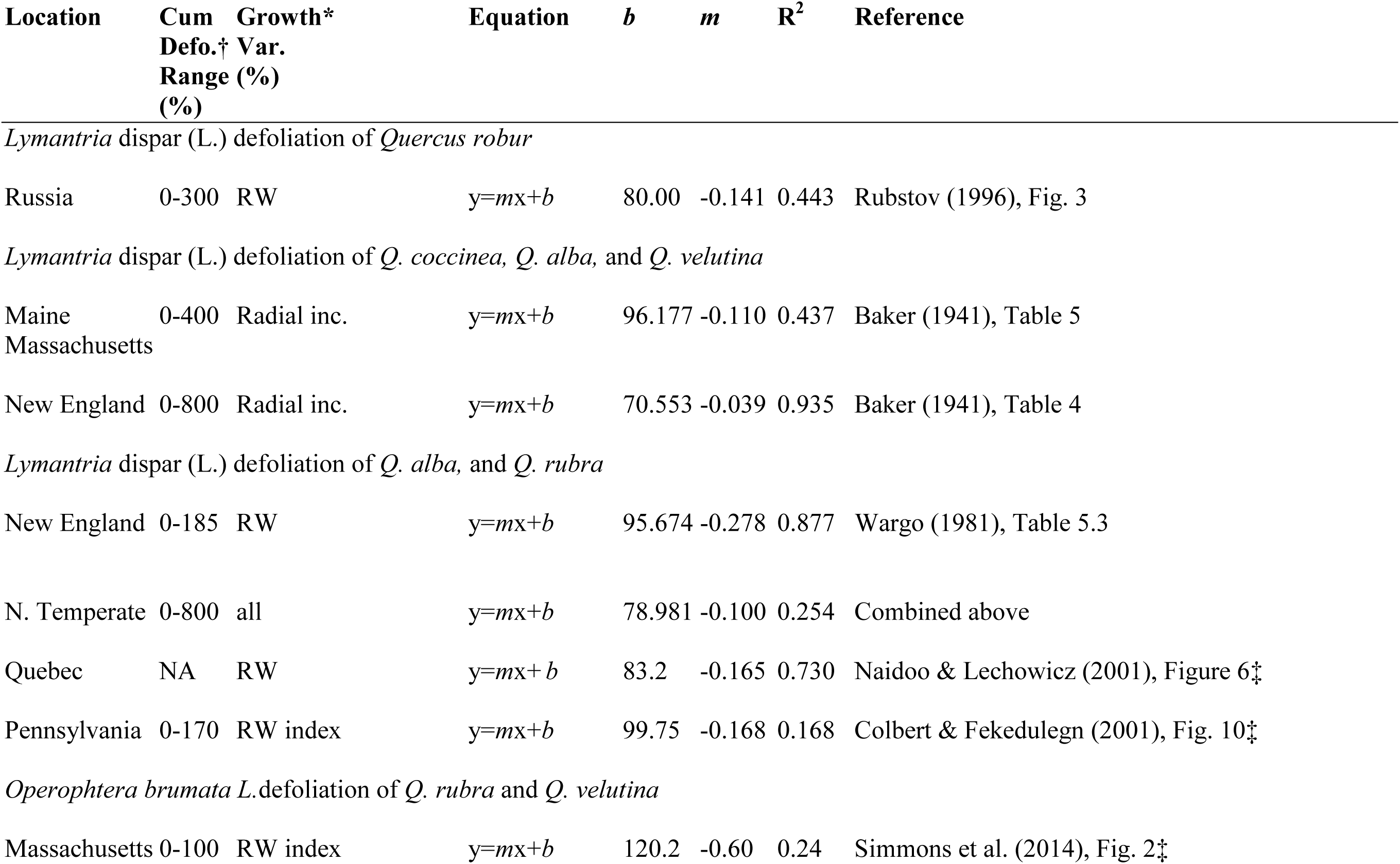
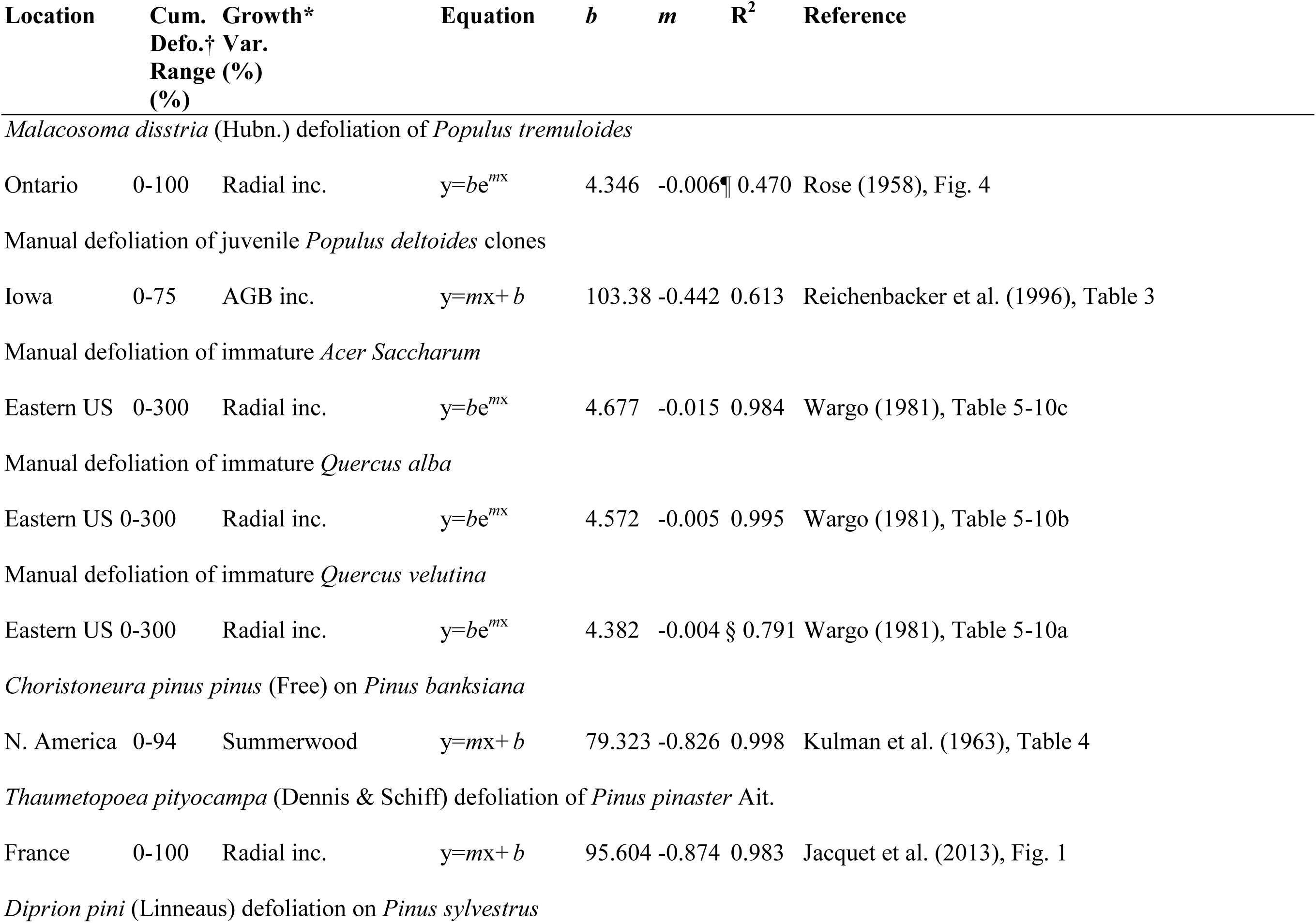
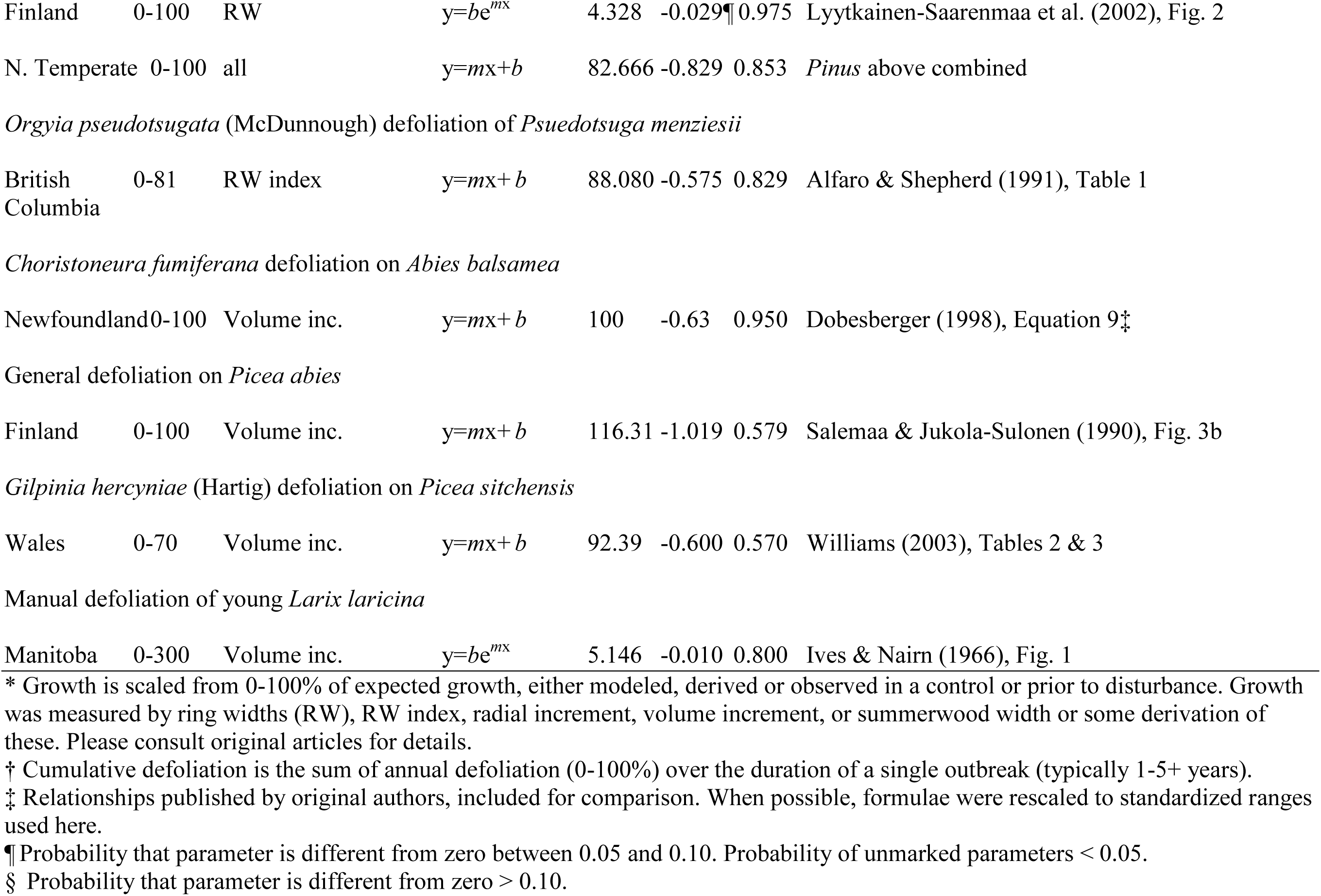
Models of relative tree growth response to cumulative defoliation. Input data were extracted and relativized from original studies to the same scale (details below).

Negative curvilinear growth responses to defoliation were mostly associated with immature trees (Fig. 2). These showed rapid early reductions in growth that were later constrained by the theoretical minimum of zero (radial wood growth cannot be negative, though stems will shrink under certain conditions (Stephens, Turner & Roo 1972; Wargo 1981; Alfaro et al. 1982; Nichols 1988; Ostaff & Maclean 1995; MacLean & MacKinnon 1996)). The curvilinear response is plausible in species that can survive many years without measurable radial growth. Moreover, linear patterns become curvilinear if they approach lower bounds to growth, and may simply represent a truncated view of the full relationship. This appears to be the case for juvenile oaks that were manually defoliated by Wargo (1981) (Fig. 2a), whose growth decreased more rapidly in a curvilinear fashion than mature, naturally defoliated oaks (Fig. 1). Manually defoliated young *L. laricina* showed a similar curvilinear response (Fig. 2a). These three patterns dominate the literature: negative curvilinear or linear growth responses with steep or gradual slopes. This suggests consistent differences in the variability and range of growth suppression among tree functional groups following defoliation stress.

### Mortality

I derived standardized cumulative defoliation and mortality data for 12 published studies covering broad areas of North American and Eurasian temperate forest, several tree species, and seven distinct spring defoliators of hardwoods, conifers or both (Table 2). Mortality increased exponentially in all examples, approaching a sigmoid curve if severe enough, or observed long enough, to saturate at 100% (MacLean 1980). This pattern was first observed by Blaise (1958) for *Abies balsamea* mortality following spruce budworm defoliation (Appendix S2). Though the available data show a consistent form for mortality relationships, the parameters varied for different tree species and defoliator systems (Fig. 3, Table 2, Appendix S2 and S3). Cumulative defoliation explained 90% of the variation in cumulative mortality of *Quercus* dominated forests following *Lymantria dispar* L. (gypsy moth (GM)) outbreaks (Table 2, Feicht et al. 1993). Mortality increased slowly at first, until cumulative defoliation reached about 150%, after which mortality increased rapidly, approaching 100% (Baker 1941; Campbell & Valentine 1972; Feicht et al. 1993; MacLean & Ebert 1999). A similar exponential increase was observed for single year defoliation levels less than or equal to 100% (Table 2, Dobbertin & Brang 2005).

**Figure 3.**
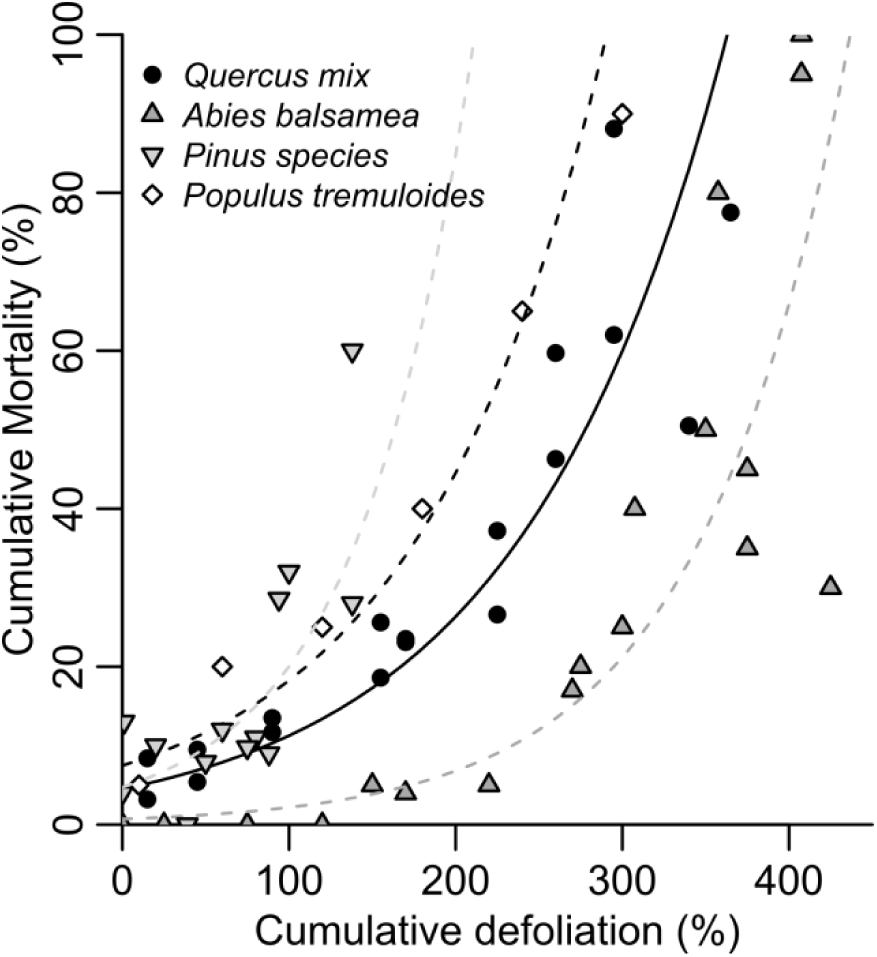
Cumulative mortality responses of example tree species to accumulated defoliation. Mortality rates rise most quickly in *Pinus* (Baker 1941, Kulman et al. 1963, Gross 1992), followed by diffuse-porous *Populus* (Mann et al. 2007), and then ring-porous *Quercus* dominated forests (Feicht et al. 1993). Mortality in *Abies* appears to rise more slowly (Blais 1958, Batzer 1973), but defoliation in this system is defined differently as damage by foliage age-class. Lines show model fits from Table 2. All plots can be found in Appendix S2.

**Table 2.**
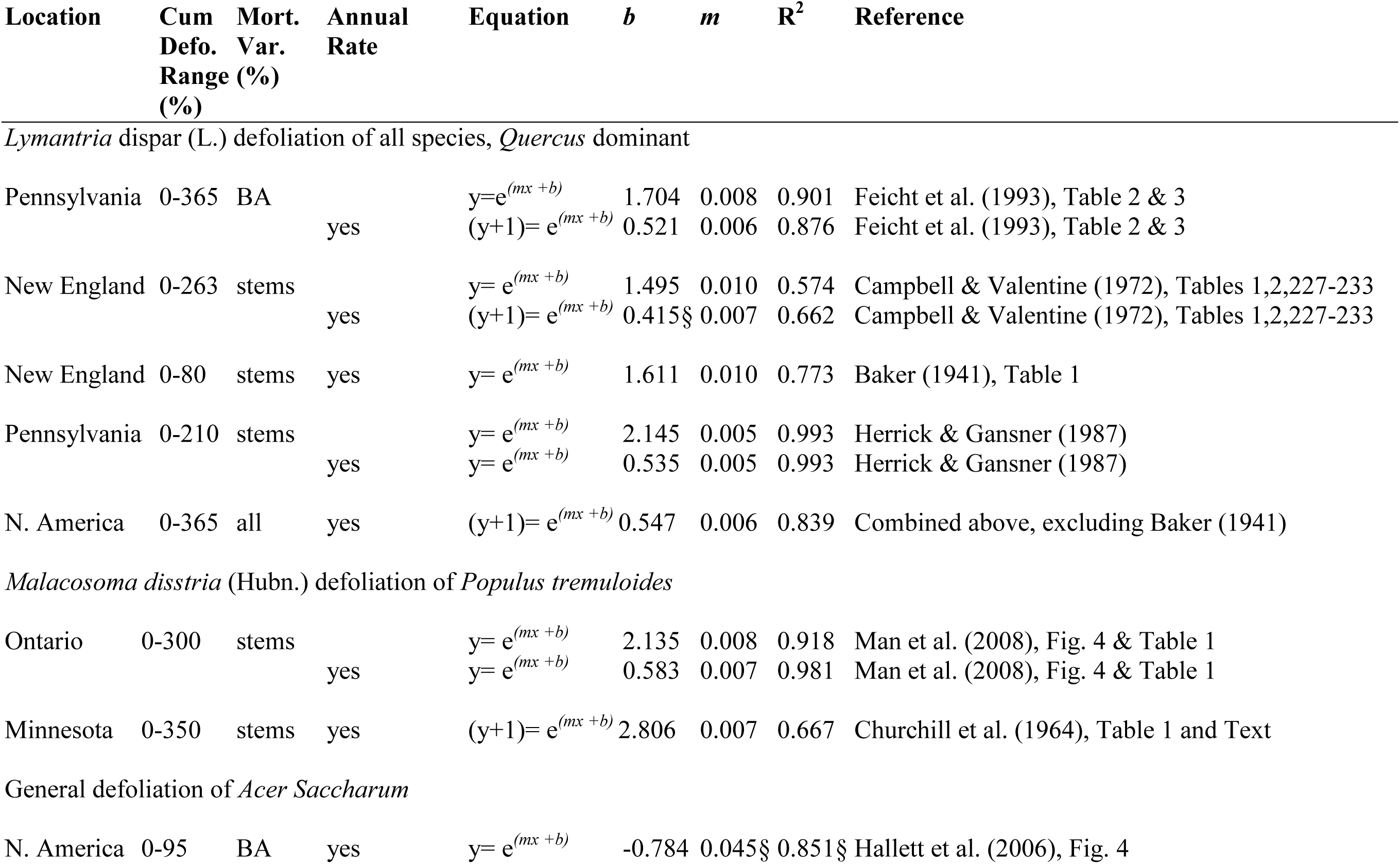
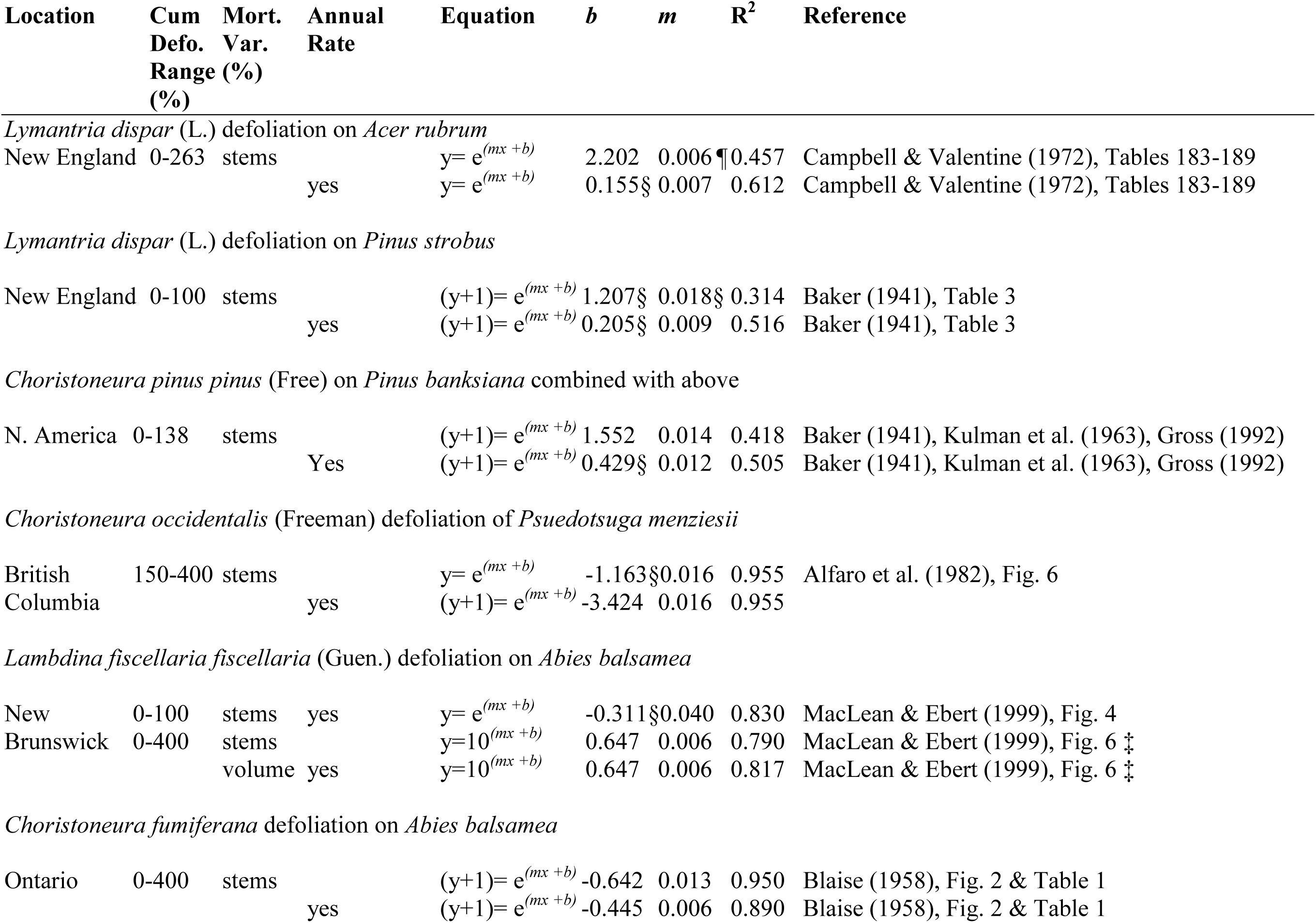
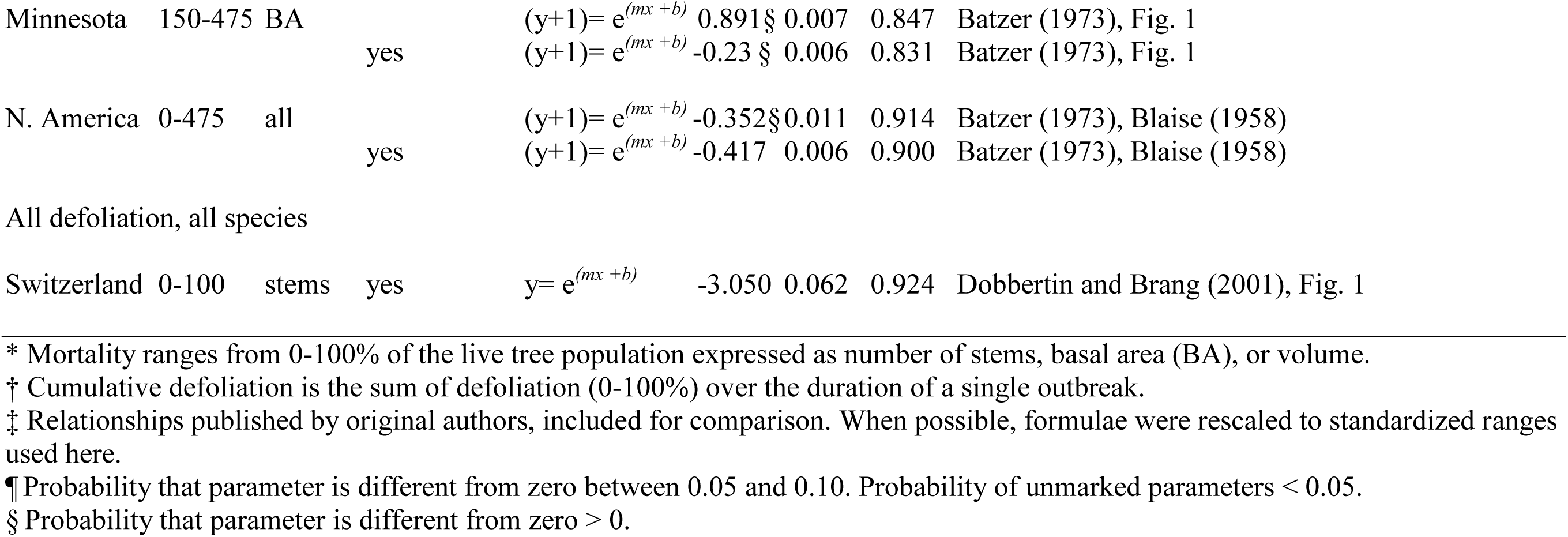
Models of cumulative tree mortality or annual mortality rates derived from cumulative defoliation.

Alfaro et al. (1982) used a powered exponential relationship to explain over 99% of the variance in mortality of Douglas fir (*Pseudotsuga mensiesii*) following a *Choristoneura occidentalis* (western spruce budworm (WSBW)) outbreak (Table 2, Appendix S2). This relationship is very similar to one derived here for *Abies balsamea* (balsam fir) mortality following *Choristoneura fumiferana* (eastern spruce budworm (SBW)) defoliation from two separate studies (Table 2, Fig. 3) (Blais 1958; Batzer 1973). These data agree with anecdotal reports of mortality becoming severe following 3-4 years of complete defoliation (Kulman 1971), though these systems take much longer than deciduous ones to reach an equivalent amount of whole canopy defoliation (as opposed to defoliation of specific foliage age-classes only). In particular, SBW outbreaks on *A. balsamea* take 10-12 years to produce equivalent damage to entire canopies because they feed preferentially on current year foliage (MacLean 1980; MacLean & Ostaff 1989). Mortality in *A. balsamea* following *Lambdina fiscellaria fiscellaria* (hemlock looper ), a defoliator of all foliage age-classes, led to a more rapid increase in mortality, closer to the range observed for *Pinus* species (Table 2, Appendix S2) (MacLean & Ebert 1999).

When data from three studies of *Pinus* response, including mortality of young *P. strobus* following GM defoliation and mature *P. banksiana* following *Choristoneura pinus pinus* (jack pine budworm (JPBW)) defoliation (Baker 1941; Kulman et al. 1963) were combined (Fig. 3), the best fit model shows mortality increasing more rapidly for defoliated *Pinus* than *Quercus* species (Fig. 3, Appendix S2 and S3). Pines are typically thought to be capable of withstanding only a few years of complete or severe defoliation, consistent with this rapid increase in mortality (O’Neil 1962; Volney 1998).

### Differences in response models: species vs. functional group

For cases where it was possible to formally compare mixed models (Appendix S3), models with species-specific parameters always ranked higher than models that varied random effects based on functional group. Species-specific random effects were best for all response models regardless of whether intercepts, slopes, or both were varied (Appendix S3). Though species-specific models represented the data best, functional group models lost little in terms of goodness-of-fit. The variance explained by the best curvilinear growth response model by functional group (R^2^c = 0.72) was not much lower than variance explained by the best species model (R^2^c=0.83) (Appendix S3) (Nakagawa & Schielzeth 2013). Variance explained by the best linear growth response models was similarly high for functional group (R^2^c=0.92) as for species (R^2^c=0.94), and the best mortality functional group model (R^2^c=0.82) lost little predictive power compared to the best species model (R^2^c=0.87). The similar predictive power of species and functional group models suggests that functional group can be used as a surrogate to predict growth and mortality response for tree species that lack data. It should be noted that mixed models that vary only intercepts (*Quercus, Pinus,* Appendix S3) may be detecting differences among the methods used by different studies to relativize growth responses against “normal” growth, rather than meaningful biological differences among species.

## Discussion

### Relationship of growth and mortality responses to phenology, wood anatomy and NSC storage

Differences in growth strategies and wood anatomy among tree functional groups may help explain the observed differences in shape and slope of the relationships reported here. For example, oaks that survive several years of gypsy moth defoliation show a linear reduction in growth of about 14-17% per 100% increase in defoliation. The best mixed models showed that slopes did not differ significantly for analogous data collected in New England, Quebec, and Russia (Baker 1941; Rubtsov 1996; Naidoo & Lechowicz 2001) (Appendix S3). Over the course of an outbreak, radial growth was reduced by an average of 50-60% in these examples, but rarely more. Temperate oaks are ring-porous species that must rely on large early wood vessels for hydraulic transport. More than 90% of these vessels cavitate under freezing winter temperatures, meaning that oaks’ hydraulic architecture will not function from one year to the next (Sperry et al. 1994; Davis, Sperry & Hacke 1999). Temperate oaks overcome this limitation by storing excess starch reserves in sapwood over winter and drawing on them to build new xylem elements the following spring (Barbaroux & Breda 2002). This early wood radial growth will actually precede flushing of the leaves by two to six weeks, and is necessary to supply water to developing foliage (Lechowicz 1984; Hacke & Sauter 1996; Suzuki, Yoda & Suzuki 1996, Michelot et al. 2012). Accordingly, many *Quercus* species, and large vessel ring-porous species in general, leaf-out later than other deciduous species (Lechowicz 1984, Panchen et al. 2014). Growth phenology research using dendrometer bands has documented examples where 30% of radial growth is completed before budburst, and 40% by the time tree canopies achieve maximum LAI in June (Kulman et al. 1963; Hinckley & Lassoie 1981; Barbaroux & Breda 2002, Zweifel et al. 2006). Similarly, 50% of the total annual xylem was formed by June for *Fraxinus excelsior,* another ring-porous species (Ladefoged 1952). As a consequence of these traits, by the time larvae of spring defoliators are large enough to cause severe defoliation damage, many oaks have already completed 30-40% of annual radial growth in the form of early wood (Kozlowski 1969, Zweifel et al. 2006).

Xylem vessels are necessary for conductance to support leaves and photosynthesis, creating a minimum cambial growth threshold for temperate, deciduous oaks that is typically 20-40% of average radial growth. The curvilinear growth response of juvenile *Quercus* trees defoliated by Wargo (1981) illustrates this well (Fig. 2a). Trees that are depleted of starch and unable to regenerate a minimum vessel layer the following spring are likely to die and leave the survivor pool. Studies of non-structural carbohydrate (NSC) storage also support this hypothesis, though the mechanisms of the carbon starvation hypothesis remain in question (Hoch, Richter & Korner 2003; Landhäusser & Lieffers 2012). Hoch et al. (2003) measured NSC allocation and reserves for seven deciduous and three evergreen tree species and estimated that typical C reserves in deciduous species could regenerate the equivalent of four canopies of foliage. This estimate agrees with anecdotal evidence that many trees die following three to four years of complete defoliation and is consistent with curvilinear mortality rates (Fig. 3, Appendix S2). Wargo (1981) measured starch reserves in juvenile *Q. velutina* and *Q. alba* as well as *Acer saccharum* following three years of 100% artificial defoliation of June foliage. Starch dropped curvilinearly to 11% and 0.2 % of average in *Q. alba* and *Q. velutina,* respectively. Starch in *A. saccharum* dropped to 0%, leading to complete mortality. Diffuse-porous species such as *A. saccharum* are less sensitive to limited radial growth. They retain multiple rings of functioning xylem to meet hydraulic requirements. They may be more sensitive to rapid reductions in starch, as they are less thrifty with NSC storage than *Quercus* species (Wargo 1981; Barbaroux & Breda 2002).

Diffuse-porous tree species from genera such as *Populus* (aspen), *Acer* (maples), and *Fagus* (beech) demonstrate a different growth phenology than oaks. This phenology could explain their wider range of radial growth response to defoliation. The smaller vessels in diffuse-porous wood rarely cavitate and water transport is thus possible using existing xylem as soon as spring conditions are suitable for budburst and shoot elongation (Lechowicz 1984; Davis et al. 1999). Diffuse-porous species typically utilize multiple annual rings for proper hydraulic function. As a result, diffuse-porous species store less starch in sapwood, only needing enough to generate a new set of foliage. These species leaf-out as early as possible when growing conditions become suitable for early leaf development (Panchen et al. 2014), and the probability of late frosts decreases (Barbaroux & Breda 2002). Diffuse-porous wood anatomy is often linked to indeterminate growth habits, which allow species to continue to grow and respond to changing conditions throughout the growing season (Kozlowski 1992). In comparison to ring-porous *Quercus*, diffuse-porous *Fagus* leaf-out earlier in the spring, do not initiate measurable radial growth until two to seven weeks after budburst has occurred, and may achieve only 5-10% of annual growth by the time canopies reach maximum LAI (Suzuki et al. 1996; Schmitt, Moller & Eckstein 2000; Barbaroux & Breda 2002, Michelot et al. 2012)). In such cases, June herbivory by defoliator larvae could suppress radial growth by up to 95%. The wider range in growth response of diffuse-porous species leads to a steeper negative slope as defoliation stress accumulates. At the same time, these strategies may improve survivability for some diffuse-porous species, such as *Populus tremuloides,* by allowing trees to dedicate more available photosynthate and NSC reserves to refoliation or chemical defense, rather than replenishing reserves for future spring wood growth. The empirical evidence in the literature remains equivocal on this topic. This review shows that defoliated young *A. saccharum* and mature *P. tremuloides* do demonstrate a wider range in radial growth suppression than *Quercus* species (Fig. 2); their growth was suppressed to below 20% of average over the observed range of defoliation.

Conifers that retain foliage for multiple years employ yet another growth strategy that contributes to a wider dynamic range of response to cumulative defoliation stress. Conifers that retain two to three years of foliage (most *Pinus*) respond differently from those that retain four to eight years of foliage age-classes (*Abies, Pseudotsuga*), as pines are generally more vulnerable to their defoliators than firs. Conductive tissue in coniferous stemwood is made up of tracheids that resist cavitation, a necessity for retention of green leaves throughout the winter dormant season (Hinckley & Lassoie 1981; Tyree & Ewers 1991). Like diffuse-porous species, conifers are not dependent on radial growth to renew xylem function and can survive periods with no radial growth. This characteristic shows up as dropped rings in dendrochronological research. Unlike deciduous trees, conifers tend to utilize older foliage more than sapwood for NSC storage, and rely on translocation from those stores to grow new foliage in the spring (Kozlowski 1992; Hoch et al. 2003). New, current-year foliage is also responsible for a greater proportion of net photosynthate production than older foliage. These characteristics mean radial growth response to defoliation can range from 0-100%, and variation in preferential herbivory of old or new foliage can produce linear or nonlinear relationships (Fig. 2b).

The simple, general relationships compiled here indicate that increasing defoliation stress slows carbon accumulation in host species through destruction of foliar biomass and suppression of radial stem growth. Productivity can be slowed at different rates, depending on the plasticity and phenology of tree growth response and defoliator characteristics. Continuous defoliation also increases tree mortality exponentially. Mortality rates increase most rapidly in *Pinus,* followed by diffuse-porous genera such as *Acer* and *Populus*, then *Quercus*, and more slowly in *Abies* and *Pseudotsuga* whose pests exhibit different feeding behavior. Nonlinear growth and mortality responses can lead to extreme short-term losses in aboveground carbon. These relationships allow us to more accurately quantify and model landscape-level effects on forest carbon accumulation. This framework and approach for quantifying the accumulation of defoliation stress and associated growth and mortality responses can be used in future empirical research to facilitate comparison across more defoliator systems.

## Acknowledgements

Funding support was provided by NASA Carbon Cycle Science grant NNX06AD45G and NASA earth system science graduate fellowship (NNX08AU93H). I thank David Mladenoff for guidance, support and editorial review, as well as B. Sturtevant, K. Raffa, M. Turner, P. Townsend, D. Lorimer, and A. Pessin for comments on early versions of this manuscript.

## Data Accessibility

– Defoliation, growth and mortality data extracted and transformed from published studies will be archived on DRYAD, as well as associated R-scripts allowing replication of the analysis.
– Appendix S1. Plots of derived growth data and back-transformed model fits by study: uploaded as online supporting information
– Appendix S2. Plots of derived mortality data and back-transformed model fits by study: uploaded as online supporting information
– Appendix S3. Model selection and parameter fits for mixed models of growth and mortality data that could be compared within defoliator systems or across species groups: uploaded as online supporting information

## References

Alfaro RI, Shepherd RF (1991) Tree-ring growth of interior douglas-fir after one year defoliation by douglas-fir tussock moth. For Sci 37: 959–964.

Alfaro RI, Vansickle GA, Thomson AJ, Wegwitz E (1982) Tree mortality and radial growth losses caused by the western spruce budworm in a douglas-fir stand in British Columbia. Can J For Res 12: 780–787.

Anderegg WRL, Callaway ES (2012) Infestation and hydraulic consequences of induced carbon starvation. Plant Physiology 159: 1866–1874.

Baker WL (1941) Effect of gypsy moth defoliation on certain forest trees. J For 39: 1017–1022.

Barbaroux C, Breda N (2002) Constrasting distribution and seasonal dynamics of carbohydrate reserves in stem wood of adult ring-porous sessile oak and diffuse-porous beech trees. Tree Physiol 22: 1201–1210.

Batzer HO (1973) Net effect of spruce budworm defoliation on mortality and growth of balsam fir. J For 71: 34–37.

Blaise JR (1958) The vulnerability of balsam fir to spruce budworm attack in northern Ontario, with special reference to physiological age of the tree. For Chron 34: 405–422.

Campbell RW, Valentine HT (1972) Tree condition and mortality following defoliation by the gypsy moth. U.S. Department of Agriculture, Forest Service, Northeastern Forest Experiment Station, Upper Darby, PA, Res. Pap. NE-236.

Churchill GB, John HH, Duncan DP, Hodson AC (1964) Long-term effects of defoliation on aspen by the forest tent caterpillar. Ecology 45: 630–636.

Colbert JJ, Fekedulegn D (2001) Effects of gypsy moth defoliation on tree growth - preliminary models for effects of cumulative defoliation on individual host tree radial increment. Proceedings: integrated management and dynamics of forest defoliating insects, Victoria, BC., U.S. Department of Agriculture, Forest Service, Northeastern Research Station

Davidson CB, Gottschalk KW, Johnson JE (1999) Tree mortality following defoliation by the European gypsy moth (*Lymantria dispar* L.) in the United States: a review. For Sci 45: 74–84.

Davis SD, Sperry JS, Hacke UG (1999) The relationship between xylem conduit diameter and cavitation caused by freezing. A J Bot 86: 1367–1372.

Dobesberger EJ (1998) Stochastic simulation of growth loss in thinned balsam fir stands defoliated by the spruce budworm in Newfoundland. Can J For Res 28: 703–710.

Feicht DL, Sandra LC (1993) Forest stand conditions after 13 years of gypsy moth infestation. Proceedings of the 9th Central Hardwood Forest Conference; Gen. Tech. Rep. NC-161, St. Paul, MN, U.S. Department of Agriculture, Forest Service, North Central Forest Experiment Station.

Foster JR, Townsend PA, DJ Mladenoff (2013). Spatial dynamics of a gypsy moth defoliation outbreak and dependence on habitat characteristics. Landscape Ecol 28: 1307–1320.

Gross HL (1992) Impact analysis for a jack pine budworm infestation in Ontario. Can J For Res 22: 818–831.

Hacke U, Sauter JJ (1996) Xylem dysfunction during winter and recovery of hydraulic conductivity in diffuse-porous and ring-porous trees. Oecologia 105: 435–439

Hallett RA, Bailey SW, Horsley SB, Long RP (2006) Influence of nutrition and stress on sugar maple at a regional scale. Can J For Res 36: 2235–2246.

Hennigar CR, MacLean DA, Norfolk CJ (2007) Effects of gypsy moth defoliation on softwood and hardwood growth and mortality in New Brunswick, Canada. Northern J Appl For 24: 138–145.

Herrick OW, Gansner DA (1987) Gypsy moth on a new frontier: forest tree defoliation and mortality. Northern J Appl For 4: 128–133.

Hinckley TM, Lassoie JP (1981) Radial growth in conifers and deciduous trees: a comparison. Mitteilungen der Forstlichen Bundes Versuchsanstalt Wien 142: 17–56

Hoch G, Richter A, Korner C (2003) Non-structural carbon compounds in temperate forest trees. Plant Cell Environ 26: 1067–1081.

Ives WG, Nairn LD (1966) Effects of defoliation on young upland tamarack in Mantioba. For Chron 42: 137–142.

Jacquet J, Orazio C, Jactel H (2012) Defoliation by processionary moth significantly reduces tree growth: a quantitative review. Ann For Sci 69: 857–866.

Jacquet J, Bosc A, O’Grady AP, Jactel H (2013) Pine growth response to processionary moth defoliation across a 40-year chronosequence. For Ecol Manage 293: 29–38.

Kozlowski TT (1969) Tree physiology and forest pests. J For 67: 118–123.

Kozlowski TT (1992) Carbohydrate sources and sinks in woody plants. Bot Rev 58: 107–222.

Kulman HM, Hodson AC, Duncan DP (1963) For Sci 9: 146–157.

Kulman HM (1971) Effects of insect defoliation on growth and mortality of trees. Annu Rev Entomol 16: 289–324.

Ladefoged K (1952) The periodicity of wood formation. Danske Videnskabernes Selskab. Biologiske skrifter. bd. 7. no. 3. København : Ejnar Munksgaard.

Landhaeusser SM, Lieffers VJ (2012) Defoliation increases risk of carbon starvation in root systems of mature aspen. Trees Struc Func 26: 653–661.

Lechowicz MJ (1984) Why do temperate deciduous trees leaf out at different times — adaptation and ecology of forest communities. Am Nat 124: 821–842.

Landhäusser SM, Lieffers VJ (2012) Defoliation increases risk of carbon starvation in root systems of mature aspen. Trees 26: 653–661.

Lyytikainen-Saarenmaa P, Tomppo E (2002) Impact of sawfly defoliation on growth of Scots pine *Pinus sylvestris* (Pinaceae) and associated economic losses. Bull Entomol Res 92: 137–140.

MacLean DA (1980) Vulnerability of spruce stands during uncontrolled spruce budworm outbreaks—a review and discussion. For Chron 56: 213–221.

MacLean DA, Ebert P (1999) The impact of hemlock looper (*Lambdina fiscerllaria fiscellaria* (Guen.)) on balsam fir and spruce in New Brunswick, Canada. For Ecol Manage 120: 77–87.

MacLean DA, Lidstone RG (1982) Defoliation by spruce budworm: estimation by ocular and shoot-count methods and variability among branches, trees, and stands. Can J For Res 12: 582–594.

MacLean DA, MacKinnon WE (1996) Accuracy of aerial sketch-mapping estimates of spruce budworm defoliation in New Brunswick. Can J For Res 26: 2099–2108.

MacLean DA, Ostaff DP (1989) Patterns of balsam fir mortality caused by an uncontrolled spruce budworm outbreak. Can J For Res 19: 1087–1095.

Man R, Kayahara GJ, Rice JA, MacDonald GB (2008) Response of trembling aspen to partial cutting and subsequent forest tent caterpillar defoliation in a boreal mixedwood stand in northeaster Ontario, Canada. Can J For Res 38: 1349–1356.

Michelot A, Simard S, Rathgeber C, Dufrene E, Damesin C (2012) Comparing the intra-annual wood formation of three European species (*Fagus sylvatica, Quercus petraea* and *Pinus sylvestris*) as related to leaf phenology and non-structural carbohydrate dynamics. Tree Phys 32: 1033–1043.

Mladenoff DJ (2004) LANDIS and forest landscape models. Ecol Modell 180: 7–19.

Naidoo R, Lechowicz MJ (2001) Effects of gypsy moth on radial growth of deciduous trees. For Sci 47: 338–348.

Nakagawa S, Schielzeth H (2013) A general and simple method for obtaining R from generalized linear mixed-effects models. Method Ecol Evol 4: 133–142.

O’Neil LC (1962) Some effects of artificial defoliation on the growth of jack pine (*Pinus banksiana* Lamb.). Can J Bot 40: 273–280.

Ostaff DP, Maclean DA (1995) Patterns of balsam fir foliar production and growth in relation to defoliation by spruce budworm. Can J For Res 25: 1128–1136.

Panchen ZA, Primack RB, Nordt B, Ellwood ER, Stevens A, Renner SS, Willis CG, Fahey R, Whittemore A, Du Y, Davis CC (2014) Leaf out times of temperate woody plants are related to phylogeny, deciduousness, growth habit and wood anatomy. New Phytol 203: 1208–1219.

Reichenbacker RR, Schultz RC, Hart ER (1996) Artificial defoliation effect on *Populus* growth, biomass production, and total nonstructural carbohydrate concentration. Popul Ecol 25: 632–642.

Rose AH (1958) The effect of defoliation on foliage production and radial growth of quaking aspen. For Sci 4: 335–342.

Salemaa M, Jukola-Sulonen E (1990) Vitality rating of *Picea abies* by defoliation class and other vigour indicators. Scand J Forest Res 5: 413–426.

Simmons MJ, Lee TD, Ducey MJ, Elkinton JS, Boettner GH, Dodds KJ (2014) Effects of invasive winter moth defoliation on tree radial growth in eastern Massachusetts, USA. Insects 5: 301–318.

Schmitt U, Moller R, Eckstein D (2000) Seasonal wood formation dynamics of beech (*Fagus sylvatica* L.) and black locust (*Robinia pseudoacacia* L.) as determined by the “pinning” technique. J Appl Bot-Angew Bot 74: 10–16.

Sevanto S, McDowell NG, Dickman LT, Pangle R, Pockman WT (2014) How do trees die? A test of the hydraulic failure and carbon starvation hypotheses. Plant Cell Environ 37: 153–161.

Sperry JS, Nichols KL, Sullivan JEM, Eastlack SE (1994) Xylem embolism in ring-porous, diffuse-porous, and coniferous trees of northern Utah and interior Alaska. Ecology 75: 1736–1752.

Stephens GR, Turner NC, De Roo HC (1972) Notes: Some effects of defoliation by gypsy moth (*Porthetria dispar* L.) and elm spanworm (*Ennomos subsignarius* Hbn.) on water balance and growth of deciduous forest trees. For Sci 18: 326–330.

Suzuki M, Yoda K, Suzuki H (1996) Phenological comparison of the onset of vessel formation between ring-porous and diffuse-porous deciduous trees in a Japanese temperate forest. Iawa J17:431–444.

Townsend PA, Singh A, Foster JR et al (2012) A general Landsat model to predict canopy defoliation in broadleaf deciduous forests. Remote Sens Environ 119: 225–265.

Tyree MT, Ewers FW (1991) The hydraulic architecture of trees and other woody-plants. New Phytol 119: 345–360.

Volney WJA (1998) Ten-year tree mortality following a jack pine outbreak in Saskatchewan. Can J For Res 28: 1784–1793.

Wiley E, Helliker B (2012) A re-evaluation of carbon storage in trees lends greater support for carbon limitation to growth. New Phytol 195: 285–289.

Williams DT, Straw NA, Day KR (2003) Defoliation of Sitka spruce by the European spruce sawfly, *Gilpinia hercyniae* (Hartig): a retrospective analysis using the needle trace method. Agr Forest Entomol 5: 235–245.

Zweifel R, Zimmermann L, Zeugin F, Newbery DM (2006) Intra-annual radial growth and water relations of trees: implications towards a growth mechanism. J Exp Botany 57: 1445–1459.

